# Investigating Neural Correlates of Emotional Regulation as a Function of Age, Race, and Socioeconomic Status

**DOI:** 10.1101/2025.08.21.671607

**Authors:** Cabria Shelton, Deanna M. Barch, Janine Bijsterbosch

## Abstract

Older adults often show improved emotional regulation with age, a phenomenon known as the aging paradox. This age-related increase in emotional regulation capacity is attributed to enhanced prefrontal cortex control over amygdala reactivity. However, because racial discrimination and economic disadvantage cause chronic stress, typical age-related neural associations may be altered in marginalized groups. Using task-functional MRI data from 8,711 UK Biobank participants aged 50-78, we investigated whether age-related associations in emotion-related brain function, specifically amygdala activation and *vmPFC–amygdala* connectivity, varied across racial and socioeconomic status (SES) groups. We found that older age was associated with decreased amygdala activation, which is consistent with improved emotional regulation. Yet, lower socioeconomic status was associated with increased amygdala activation, suggesting heightened stress-related reactivity. No significant age-related effects on *vmPFC–amygdala* connectivity were observed at the population level. Black participants showed a stronger age-related decline in functional connectivity compared to other racial groups. These findings call for more inclusive and diverse neuroimaging studies to better understand brain health across marginalized groups.

## Introduction

Studies suggest that emotional regulation often improves across the adult lifespan (Prakash et al., 2014), despite declines in cognitive control with age. Both emotional regulation and cognitive control processes rely on dynamic interactions between the prefrontal cortex (PFC) and the amygdala (Ray & Zald, 2012). However, task-based functional MRI (fMRI) research on age-related emotional regulation has yielded inconsistent behavioral findings. Some studies report a decline in the ability to down-regulate negative affect (Schweizer et al., 2019) while others report variability in amygdala-prefrontal effective connectivity during reappraisal (Morawetz et al., 2017), making it challenging to draw consistent conclusions. These inconsistencies highlight the need for more nuanced investigations into the neural mechanisms underlying emotional regulation with age.

Although the prefrontal cortex and amygdala have been consistently implicated in emotion processing and regulation tasks, little is known about how their activity varies across different social and demographic contexts (Dhamala et al., 2025). To date, many neuroimaging studies on emotional regulation have predominantly consisted of white, high socioeconomic status (SES) samples (Ricard et al., 2023). As a result, the lack of diversity limits generalizability and obscures potential differences in brain-behavior relationships across marginalized populations. For instance, Ricard et al. 2023 noted that brain-behavior associations from datasets dominated by white Americans may fail to generalize to underrepresented groups. Similarly, Thalmayer et al. 2021 emphasized that low diversity in data collection undermines generalizability and relevance to the broader population. Thus, previous findings on emotional regulation in homogenous samples (Ricard et al., 2023; Thalmayer et al., 2021) do not tell us whether typical age-related patterns in emotional regulation may differ in marginalized groups.

Lifelong exposure to structural inequalities, including racial discrimination and economic hardship, has been associated with altered brain function, particularly in brain regions associated with affective processing (Webb et al., 2024). Previous studies have linked this chronic stress to increased amygdala activation, reduced amygdala volume, and disrupted PFC activity (Fiocco, 2024). These neural differences related to adversity could moderate the typical associations of emotional regulation in aging. Therefore, understanding how emotional regulation unfolds cross-sectionally with age in marginalized groups would benefit from integrating social determinants into neuroimaging analyses.

The present study aims to investigate how age-related differences in emotion-related brain function vary by race and SES. We focus on two brain regions critical to emotional regulation: the amygdala and ventromedial prefrontal cortex (vmPFC). The amygdala plays a central role in processing emotional stimuli, fear, and anxiety (Phelps & LeDoux, 2005; Šimić et al., 2021), and the vmPFC is involved in top-down regulation of stress responses (Suzuki & Tanaka, 2021). A recent study suggests that the vmPFC is involved in increasing stress coping when individuals are exposed to stressful images (Sinha et al., 2016), highlighting the importance of the vmPFC in emotional regulation, potentially via regulation of amygdala activity. In this study, we examined whether age-related patterns of amygdala activation and vmPFC–amygdala functional connectivity differed across racial and socioeconomic groups.

To address this aim, we utilized data from the UK Biobank, a large-scale cohort that includes neuroimaging and biomedical data. We focused on task-based functional MRI (tfMRI) data collected during the Hariri Faces Emotion Task, where participants matched emotionally charged (angry or fearful) expressions or neutral geometric shapes. This task reliably engages the amygdala and prefrontal cortex (Hariri et al., 2002; Preckel et al., 2019), offering robust insight into neural processes involved in emotion processing and regulation (Banks et al., 2007). By analyzing these neural markers across different racial and SES groups, we aimed to clarify how marginalization alters age-related patterns in emotional regulation.

## Methods & Materials

We conducted a cross-sectional analysis of UK Biobank tfMRI data from 8,711 participants aged 50 to 78 years (mean age of 62.82 years; 52.5% female) with complete race information and covariates at a single imaging visit. Variable descriptions and UK Biobank field IDs are provided in Table 1.

**Table 1.** UK Biobank Variables Used.

| <u>Variable Description</u> | <u>UKB Field ID</u> | <u>Notes</u> | <u>Used in Models As</u> |
| --- | --- | --- | --- |
| Townsend Deprivation Index | 22189-0.0 | Higher values = more deprivation | Predictor (Socioeconomic Status ) |
| Race/Ethnicity | 21000-2.0 | Categorical- (e.g, White-British, Asian-Indian, Black-Caribbean, etc.) | Predictor |
| vmPFC–amygdala FC | N.A. | Fisher Z -Z-transformed | Outcome |

|  |  | Pearson correlation |  |
| --- | --- | --- | --- |
| Amygala Activation<br>(Faces > Shapes) | 25054-2.0 | tfMRI Median<br>z-statistic in<br>group-defined mask | Outcome |
| Age | 21003-2.0 | Continuous, in years | Covariate (control<br>for confounds) |
| Sex | 31-0.0 | Coded 0 = female, 1<br>= male | Covariate (control<br>for confounds) |
| Site | 54-2.0 | Categorical, imaging<br>center | Covariate (control<br>for confounds) |
| Intracranial Volume<br>(ICV) | 26521-2.0 | Continuous | Covariate (control<br>for confounds) |
| Mean tfMRI Head<br>Motion | 25742-2.0 | Mean motion during<br>the Hariri task,<br>averaged across<br>space and time<br>points | Covariate (control<br>for confounds) |

To ensure consistency and comparability across models, we restricted analyses to participants with complete race data during the imaging visit. For SES models, the sample included 8,706 participants. Race-stratified analyses included 8,711 participants (White: 8,483; Black: 52; Asian: 103) and a slightly smaller sample for functional connectivity (FC), the final (N = 8,611; White: 8,385; Black: 51; Asian: 102), reflecting a minor amount of missing connectivity data.

### Task fMRI Preprocessing

All UKB tfMRI data were obtained using the 3T Siemens Skyra scanner and preprocessed using the pipeline described by Alfaro-Almagro et al., 2018 based on the FMRIB Software Library. UKB brain imaging modalities include T1-weighted structural (T1), T2-weighted FLAIR imaging (T2), Susceptibility-weighted imaging (swMRI), diffusion MRI (dMRI), and functional modalities: resting-state fMRI (rsfMRI) and task-based fMRI (tfMRI). According to the UKB Brain Imaging Documentation (https://biobank.ctsu.ox.ac.uk/crystal/crystal/docs/brain_mri.pdf), the tfMRI task was a minute version of the Hariri faces/shapes emotions task, originally used in the Human Connectome Project (Hariri et al., 2002). Participants were presented with angry or fearful expressions or shapes and asked to match the top picture with the corresponding bottom picture.

### Imaging Phenotypes

All imaging preprocessing and regions of interest (ROI) based analysis were performed using the FMRIB Software Library (FSL, v6.07.8; see Figure 1). Standard space ROI masks were nonlinearly warped to subject-specific functional space using applywarp, with the reference image MNI152_T1_2mm_brain.nii.gz. Time series were extracted using fslmeants.

**Figure 1.**
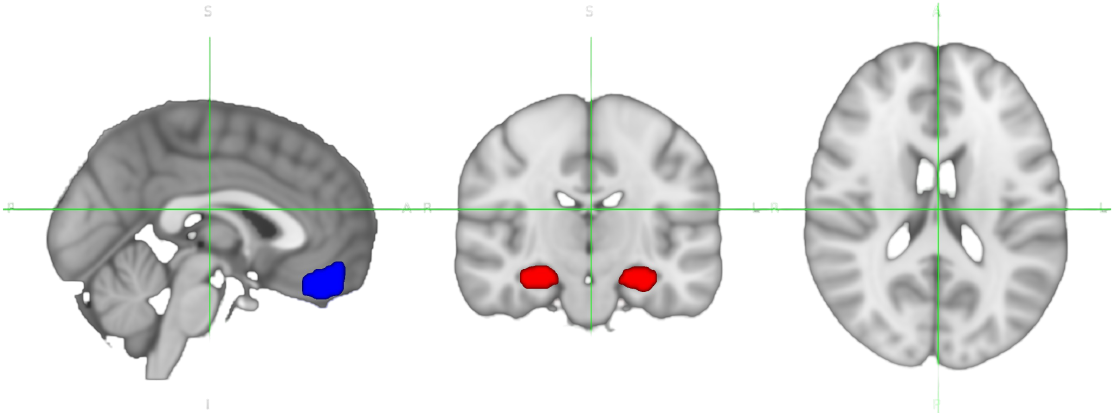
Regions of interest (ROIs) used in the analysis. Amygdala and ventromedial prefrontal cortex (vmPFC) ROIs were defined using the Harvard-Oxford probabilistic atlas and nonlinearly warped to subject-specific functional space. Time series were extracted from these ROIs to compute task-based activation and functional connectivity metrics.

Following image preprocessing, we extract two imaging-derived phenotypes of interest as described below.

1. Amygdala activation, defined as the median z-statistic from the faces > shapes contrast within a group-defined amygdala activation mask (UKB Field 25054). The z-statistics were estimated using a general linear model (GLM) fit with autocorrelation correction as implemented in the FSL’s FEAT, modeling task conditions across the full time series to quantify voxelwise activation^4^.
2. Amygdala-ventromedial prefrontal cortex connectivity (vmPFC), computed as the Fisher z-transformed Pearson correlation between time series extracted from the amygdala and vmPFC ROIs. Both ROIs were defined using thresholded and binarized masks from the Harvard-Oxford probabilistic atlas: combining left and right amygdala for the amygdala mask, and using the frontal medial cortex region for the vmPFC mask. Masks were nonlinearly warped to each subject’s functional space using FSL’s applywarp, with MNI152_T1_2mm_brain.nii.gz as the reference image. Time series were extracted using fslmeants and were not residualized for task activation effects, so the resulting connectivity reflects the coactivation between regions during task performance.

### Analyses

We used ordinary least squares (OLS) linear regression to test associations between age and two brain phenotypes described above. All variables included in the analyses were z-scored prior to modeling to ensure standardized interpretation of regression coefficients. All models included the following covariates: sex, imaging site, intracranial volume, and mean tfMRI head motion (see Table 1 for field IDs). Separate models were developed to assess the impact of socioeconomic status (Townsend Deprivation Index) and race, as described below.

### Socioeconomic Status (SES) Models

Socioeconomic status was operationalized using the Townsend Deprivation Index, with higher values indicating greater deprivation. We examined whether the relationship between age and each brain phenotype varied by SES using the following linear regression model:

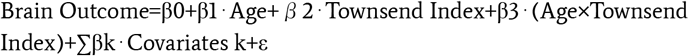

The model was applied separately to amygdala activation and functional connectivity outcomes.

### Race Stratified Models

Race was self-reported and categorized as White, Black, or Asian. There were no Hispanic or Latino participants present in the sample. Due to the categorical nature of race, regression analyses against age were performed separately within each category. Within each racial group, we applied the following model separately for amygdala activation and vmPFC–amygdala connectivity outcomes:

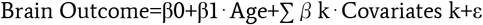

Due to large sample size differences between racial groups, the resulting beta values for age cannot be directly compared to assess racial differences. To enable fair statistical comparisons, we used a bootstrapping approach. Specifically, we drew 2,000 random subsamples samples of White participants matched in size to the total available Asian (N=103 for activation and N=102 for connectivity) and Black (N=52 for activation and N=51 for connectivity) groups. Within each subsample, we re-estimated the regression model to generate distributions of age beta values for age from equally sized White groups.

To evaluate racial differences in age-related effects, we subsequently compared the observed age coefficient from the Black and Asian models to the distribution of age coefficients obtained from the White bootstrap samples. Based on prior evidence that chronic stress exposures in marganilized groups are more likely to amplify age-related declines, we expected any group difference to occur in a single direction (i,e., a more negative age effect). For this reason, we report one-sided bootstrap p-values, calculated as the proportion of White subsamples with age coefficients equal to or more negative than the observed coefficient in the comparison group.

## Results

### Preliminary Analyses

We first examined correlations among key variables to check for multicollinearity and assess their relationships. Figure 2 shows the correlation matrix for brain outcomes, key predictors, and covariates used throughout all analyses.

**Figure 2:**
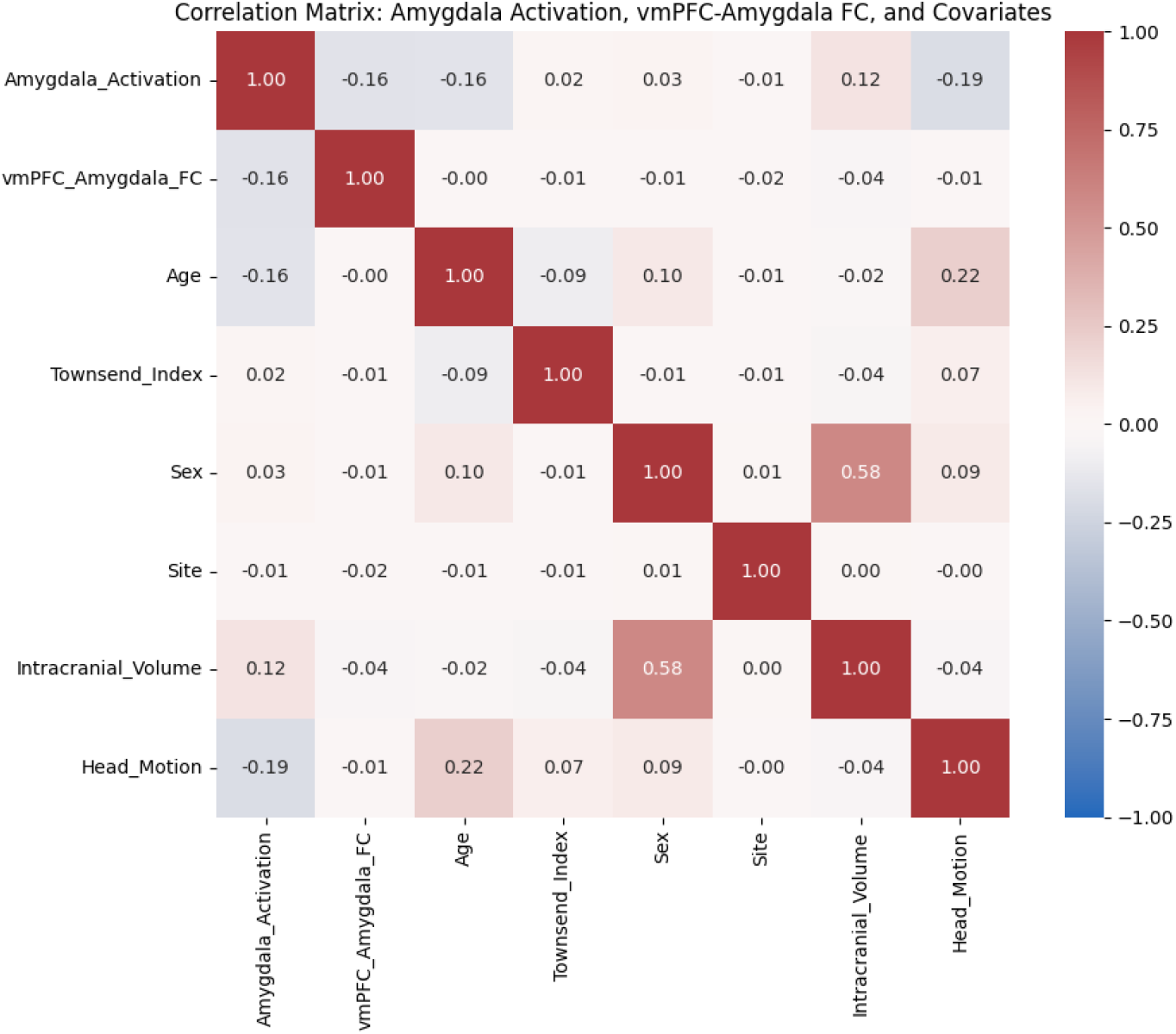
Correlation matrix showing Pearson’s correlation coefficients among brain outcomes, predictors, and covariates. Values are color-coded (red = positive, blue = negative). Strongest relationships are observed between sex and intracranial volume, and between head motion and amygdala activation.

No correlations exceeded r = 0.60, indicating that none of the variables were too highly correlated and thus multicollinearity is unlikely to cause bias in regression estimates, especially because the highest correlations were observed between covariates of no interest. The highest correlation among covariates was between sex and ICV. vmPFC-amygdala functional connectivity was not highly correlated with any predictors or covariates. Amygdala activation showed negative correlations with age (r = -0.16), head motion (r = -0.19), and a slightly positive correlation with ICV (r = 0.12). Prior evidence suggests that emotional regulation improves with age. Our finding of a negative correlation (r = -0.16) between age and amygdala activation in the full sample is consistent with such prior findings.

Race was not included in the correlations described above due to its categorical nature. To examine the association between race and socioeconomic status, we plotted the distribution of the Townsend index as a function of race. The results reveal high collinearity between race and socioeconomic status, with systematically higher Townsend index (i.e., lower SES) for Black participants compared to White and Asian participants (Figure 3). As such, we performed separate analyses to assess the role of race and SES below (instead of a combined model), to avoid collinearity.

**Figure 3:**
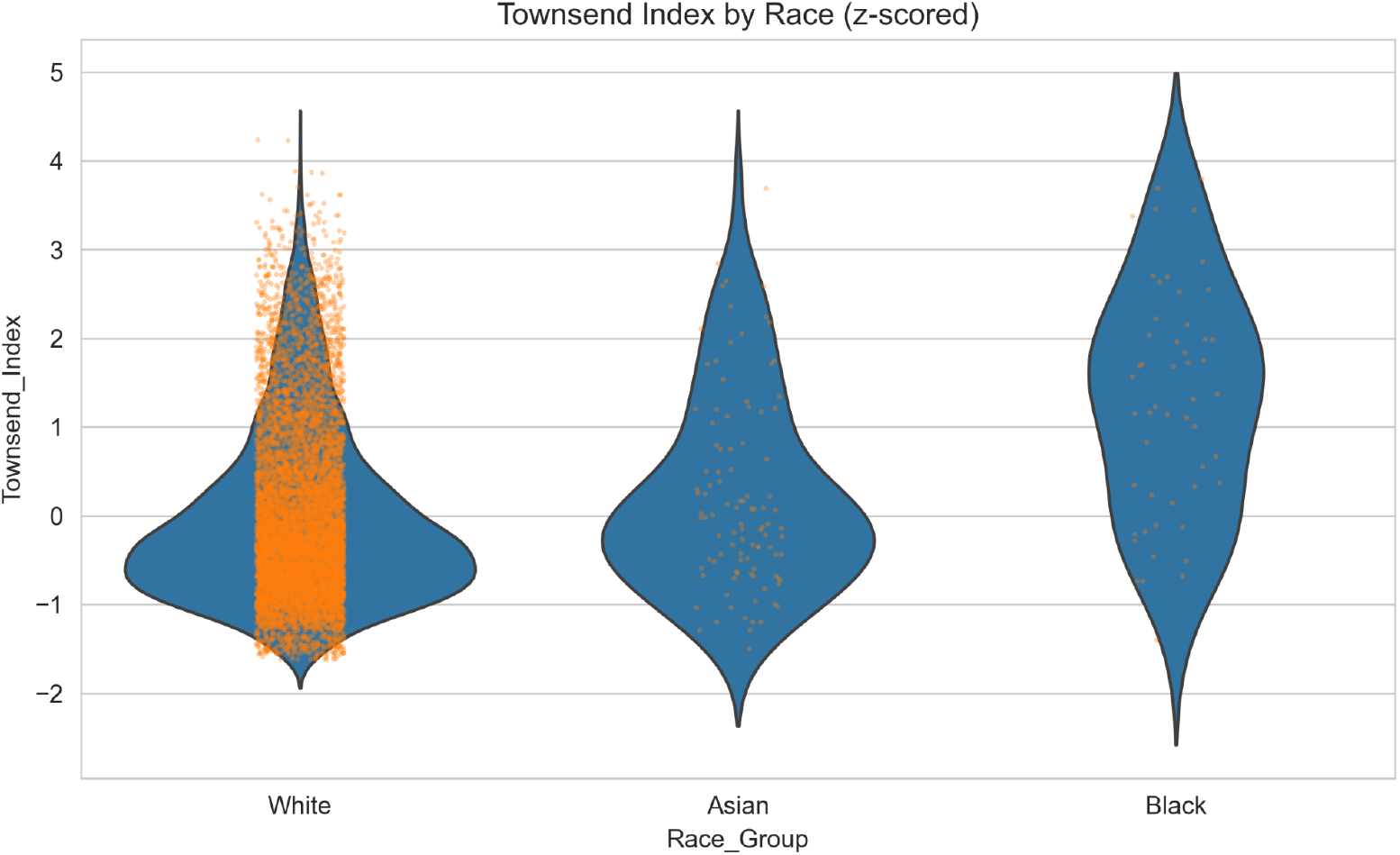
Townsend Deprivation Index by Racial Group. Violin plots illustrating the distribution of z-scored Townsend Deprivation Index values within each racial group; orange dots represent individual participants. Higher values indicate greater socioeconomic deprivation and, therefore, lower SES status. Black participants exhibit higher deprivation on average than White and Asian participants.

### Socioeconomic Differences in Age-Related Neural Associations

#### Amygdala Activation

Older age was significantly associated with reduced amygdala activation (*β* = -0.094, p < 1e-70; Figure 4). Socioeconomic status, indexed by the Townsend Deprivation Index, was also a significant predictor, with lower SES (i.e, higher deprivation scores) associated with increased amygdala activation (*β* = 0.013, p = 0.008). However, the age x SES interaction term was not statistically significant (*β* = 0.002, p = 0.668), providing little evidence that SES moderates age in relationship to amygdala activation.

**Figure 4:**
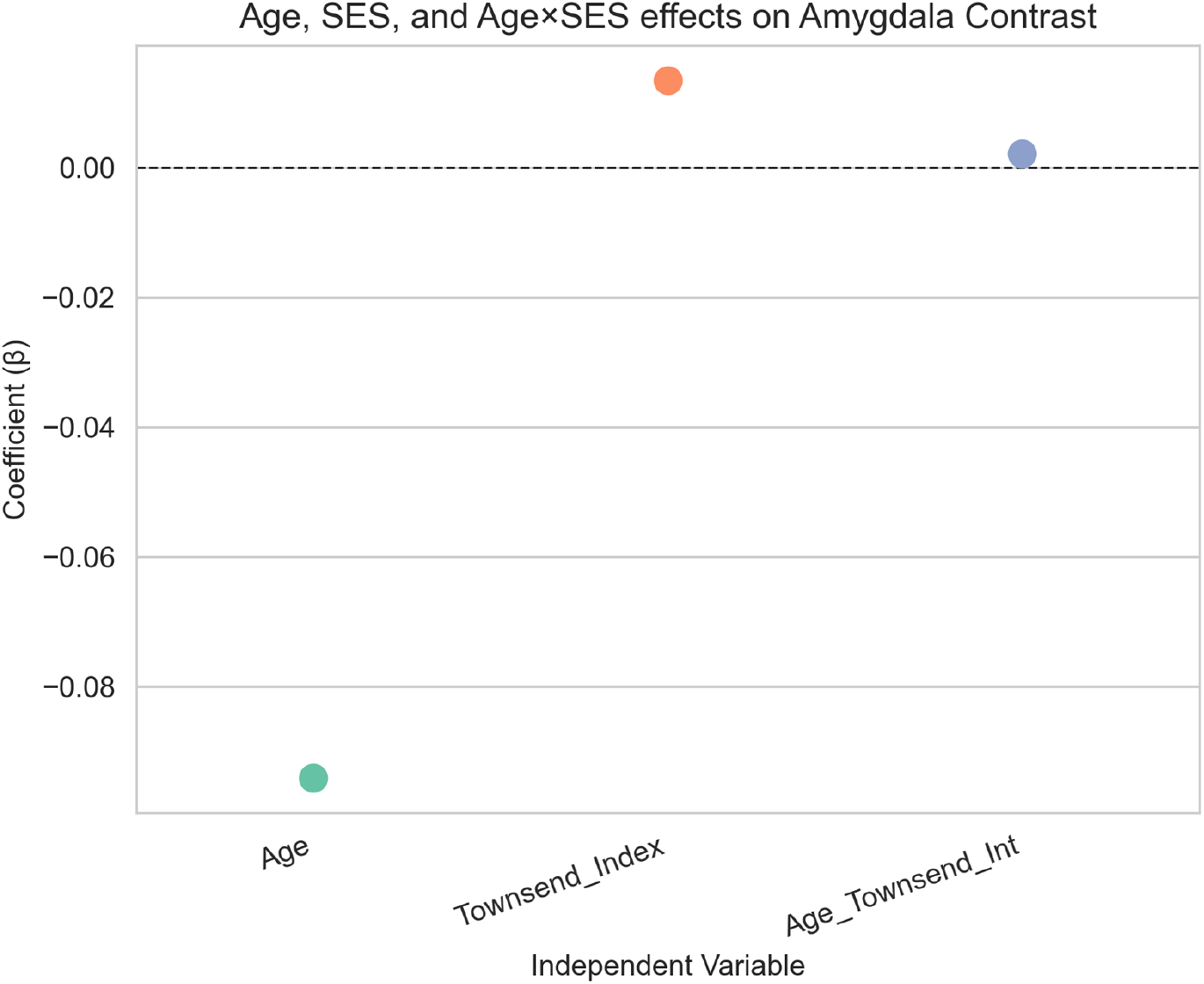
Amygdala Activation – SES Model. Association between age, socioeconomic status (SES), and amygdala activation. Older age was significantly associated with reduced amygdala activation, while greater deprivation (higher Townsend Index scores) was associated with increased amygdala activation. No significant age × SES interaction was observed.

#### vmPFC–amygdala Functional Connectivity

SES was not significantly associated with *vmPFC–amygdala* connectivity (*β* = -0.008, p = 0.540). Neither age (*β* = -0.007, p = 0.540) nor the age x SES interaction term (*β* = 0.015, p = 0.158) was significant (Figure 5), suggesting limited evidence that SES affects age-related patterns of functional connectivity.

**Figure 5:**
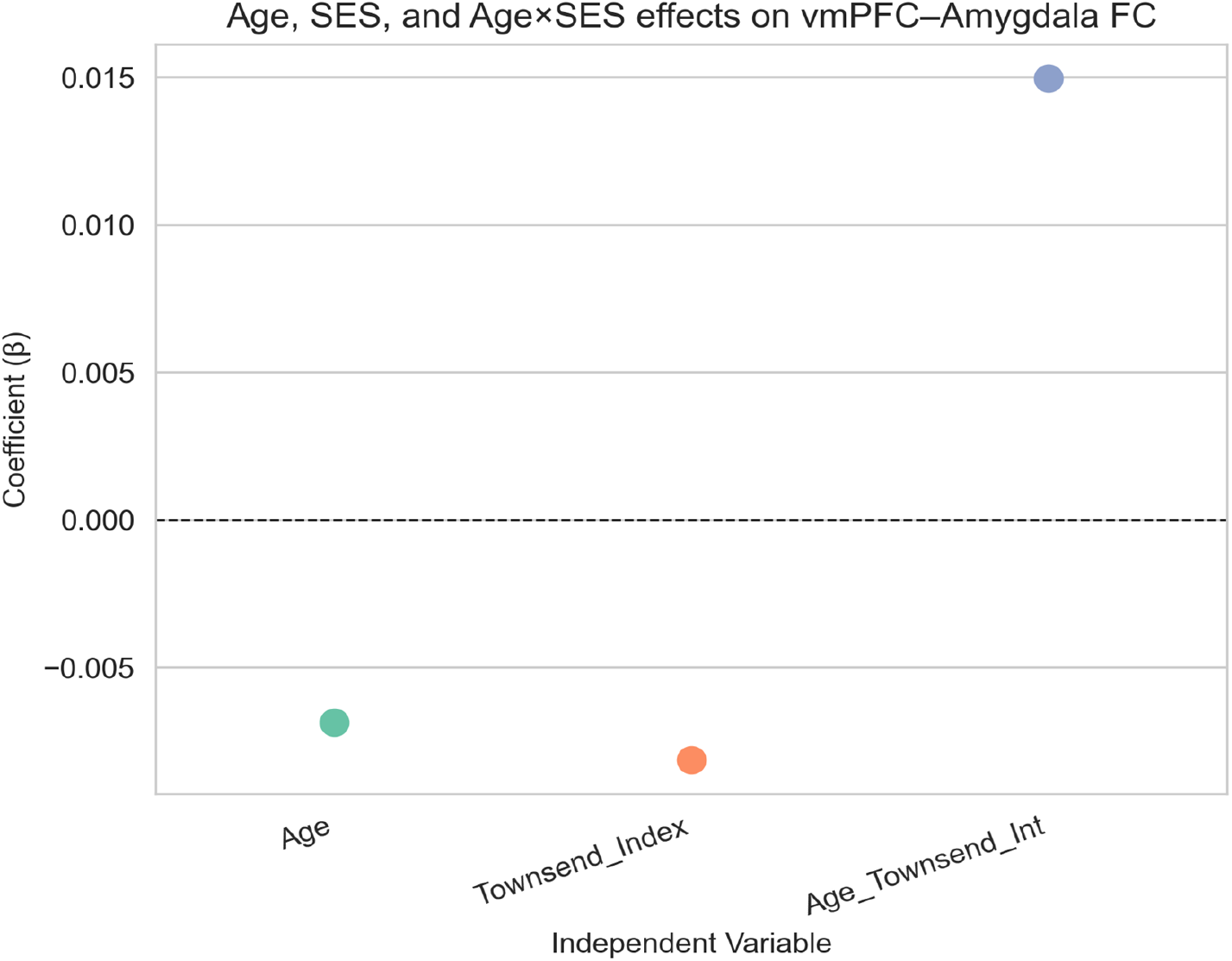
vmPFC–Amygdala Connectivity – SES Model. Association between age, socioeconomic status (SES), and amygdala–vmPFC functional connectivity. No significant effects of age, SES, or their interaction were observed on functional connectivity between the amygdala and vmPFC.

### Racial Differences in Age-Related Neural Associations

#### Amygdala Activation

Age was significantly associated with reduced amygdala activation among White (p =5.11e-27) and Asian (p = 0.016) participants, but this effect was not observed among Black participants (p = 0.46; Figure 6).

**Figure 6:**
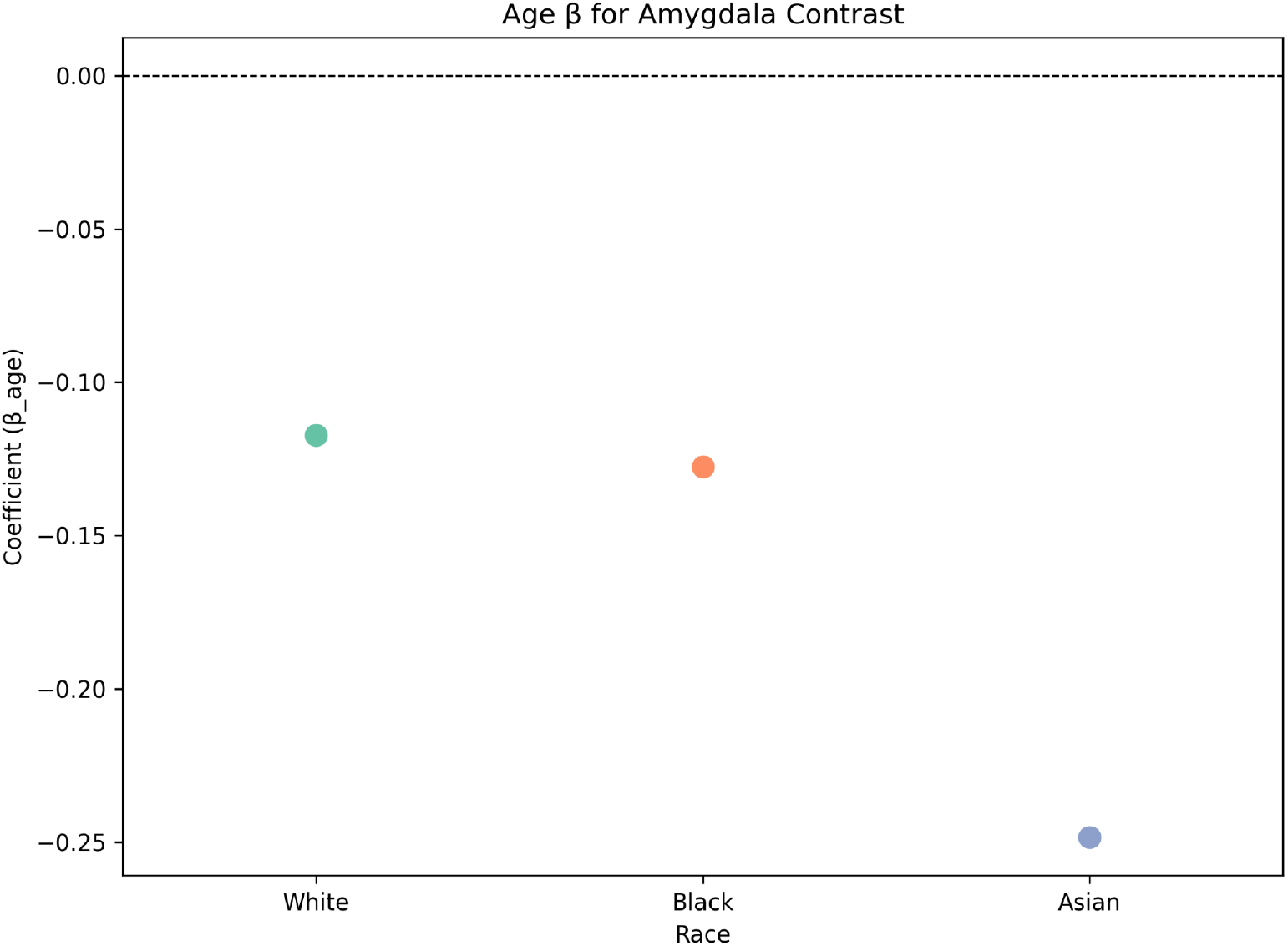
Linear regression coefficients for age-related effects on amygdala activation across racial groups. Negative coefficients indicate lower amygdala activation with age (the age x race interaction was not included due to race being a categorical variable).

Given the considerable imbalance in subgroup sizes (White: 8,483; Black: 52; Asian: 103), we then performed sample size-matched bootstrapping to enable valid statistical comparisons of associations between age and amygdala activation across racial groups. Bootstrap comparisons showed no significant differences in age-related effects for the Black (Figure 7: p> 0.05) or Asian participants (Figure 8; p > 0.05) relative to sample size-matched White results, suggesting observed racial group differences in amygdala activation in Figure 6 may be largely due to power differences rather than true age-related neural associations.

**Figure 7:**
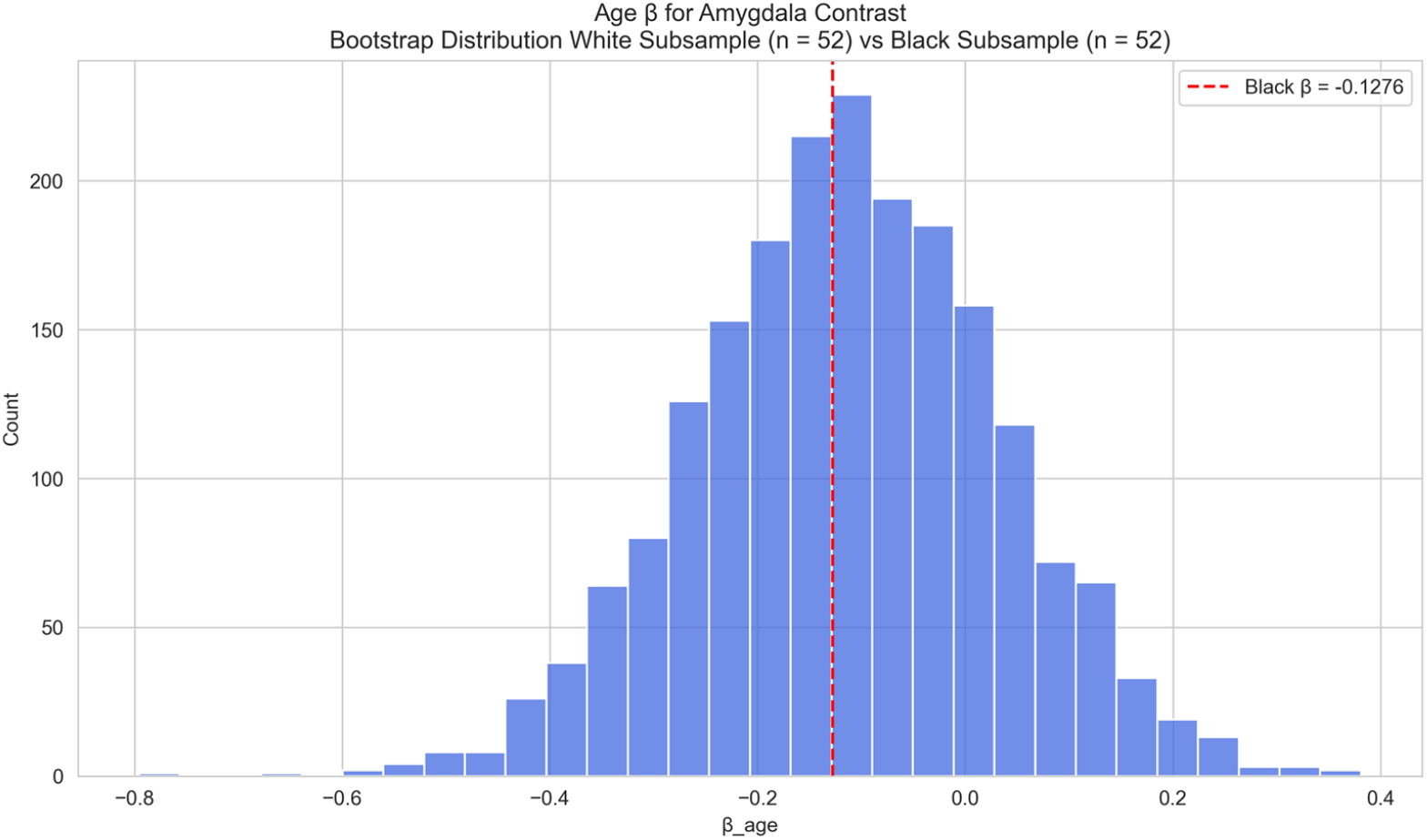
Amygdala Activation – Race-Stratified (Black vs White) Race-stratified analysis of age effects on amygdala activation: Black versus White participants. The observed age-related coefficient for the Black subgroup is shown as a vertical red line. Age was associated with reduced amygdala activation in White participants but not in Black participants. Bootstrap comparisons did not reveal significant differences in age-related effects between groups.

**Figure 8:**
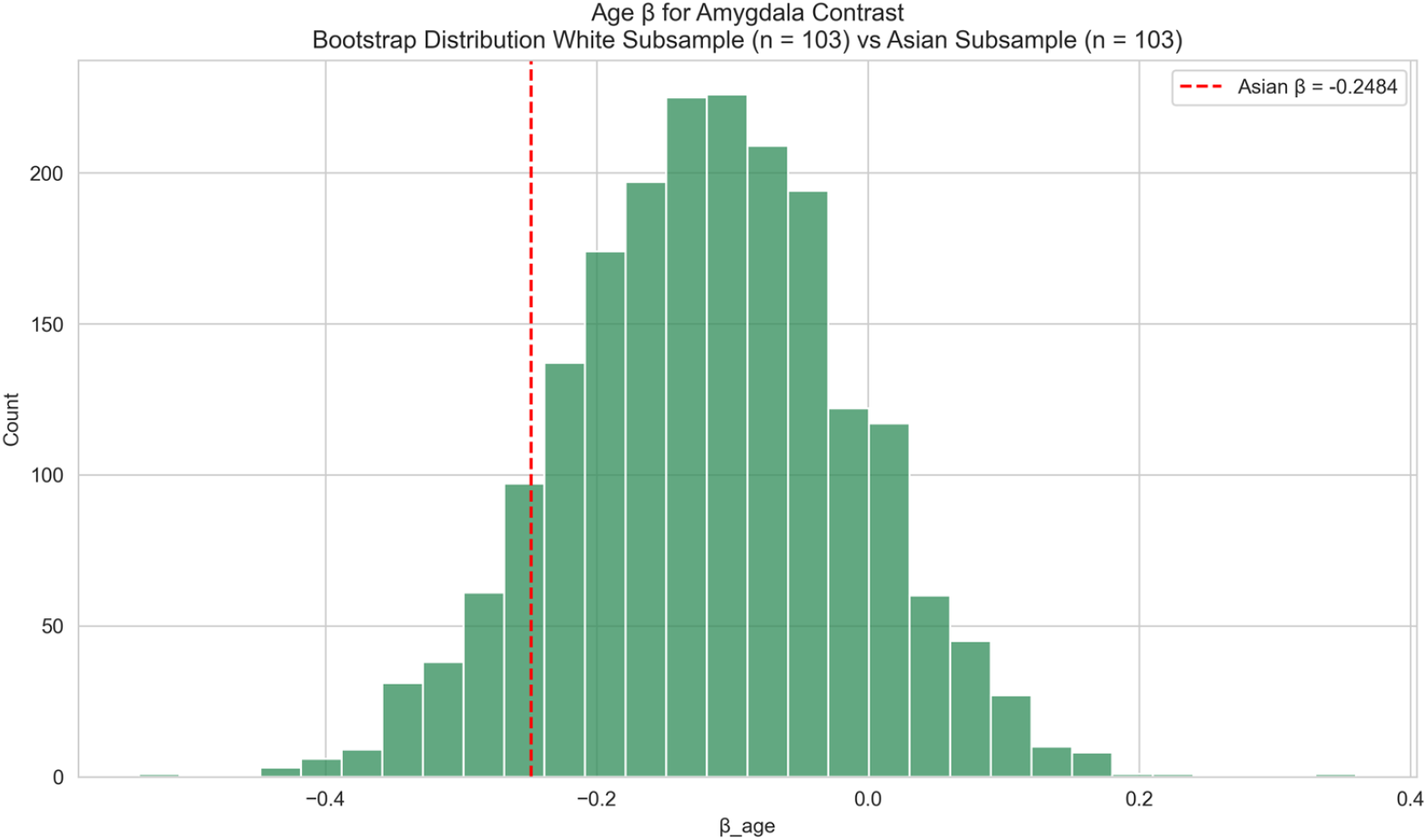
Amygdala Activation – Race-Stratified (Asian vs White. Race-stratified analysis of age effects on amygdala activation: Asian versus White participants. The observed age-related coefficient for the Asian subgroup is shown as a vertical red line. Age was associated with reduced amygdala activation in Asian participants. Bootstrap comparisons showed no significant difference in age-related effects between Asian and White participants.

#### vmPFC-amygdala Functional Connectivity

For vmPFC-amygdala connectivity, no significant age-related effects were observed for White (β = -0.0057, n=8,385) or Asian (β = -0.0291, n=102) participants. In contrast, Black participants showed a stronger negative age-related effect (β = -0.3063, n = 51). Figure 9 shows these differences via swarm plot visualization, emphasizing the notably larger negative coefficient for the Black subgroup compared to the White and Asian subgroups.

**Figure 9:**
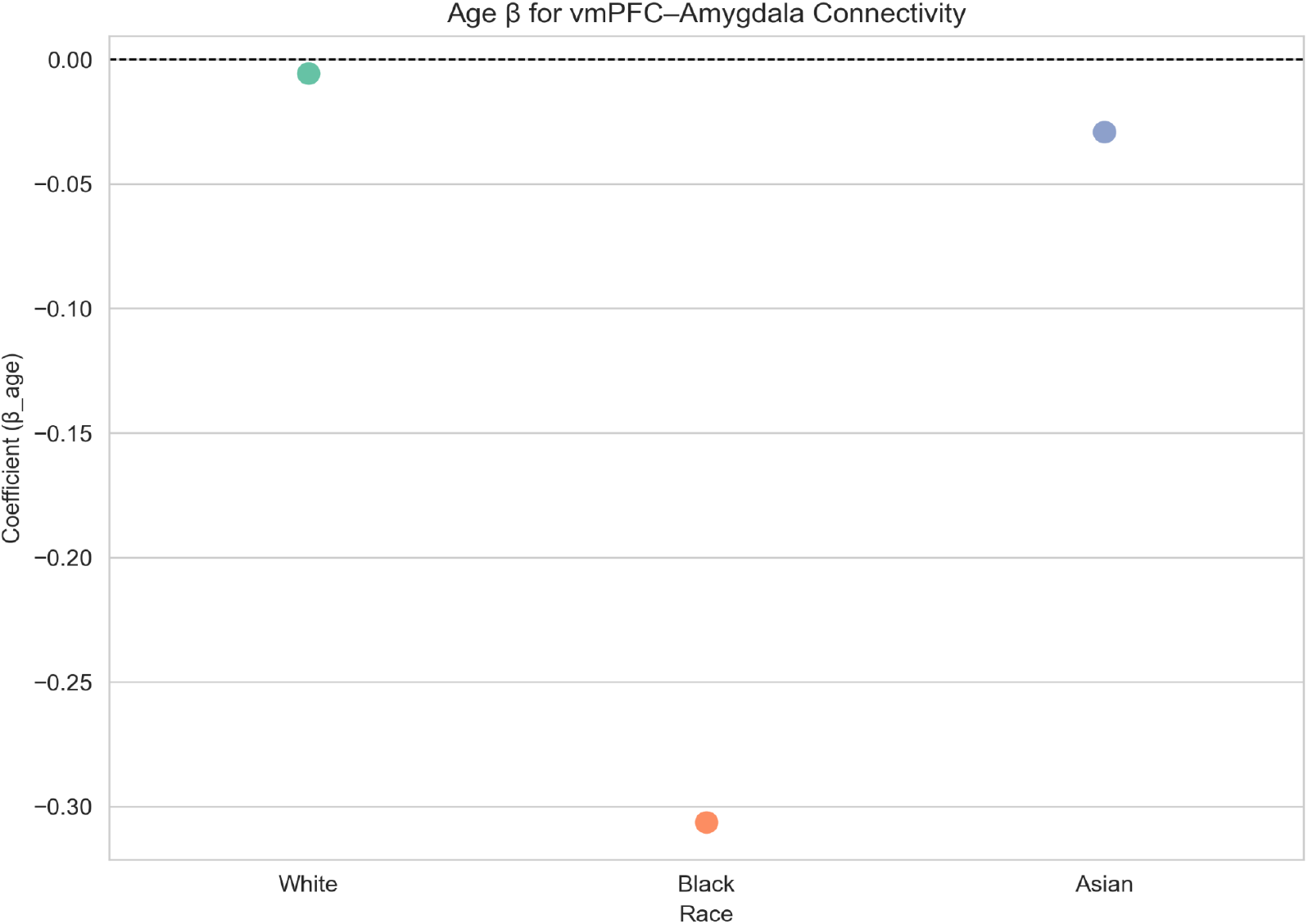
Linear regression coefficients for age-related effects on vmPFC-amygdala functional connectivity across racial groups. Negative coefficients indicate lower amygdala activation with age (the age x race interaction was not included due to race being a categorical variable).

Again, given the small Black subgroup size, bootstrap analyses were conducted to determine if this stronger effect reflected a meaningful difference or was simply due to sample size differences. The bootstrap analysis (matched White samples, Figure 10) indicated the negative age-effect in Black participants was significantly greater (one-sided p = 0.027), suggesting a significant racial difference in age-related decline of vmPFC-amygdala FC. No significant difference was found between Asian participants and matched White samples (Figure 11; p=0.3885).

**Figure 10:**
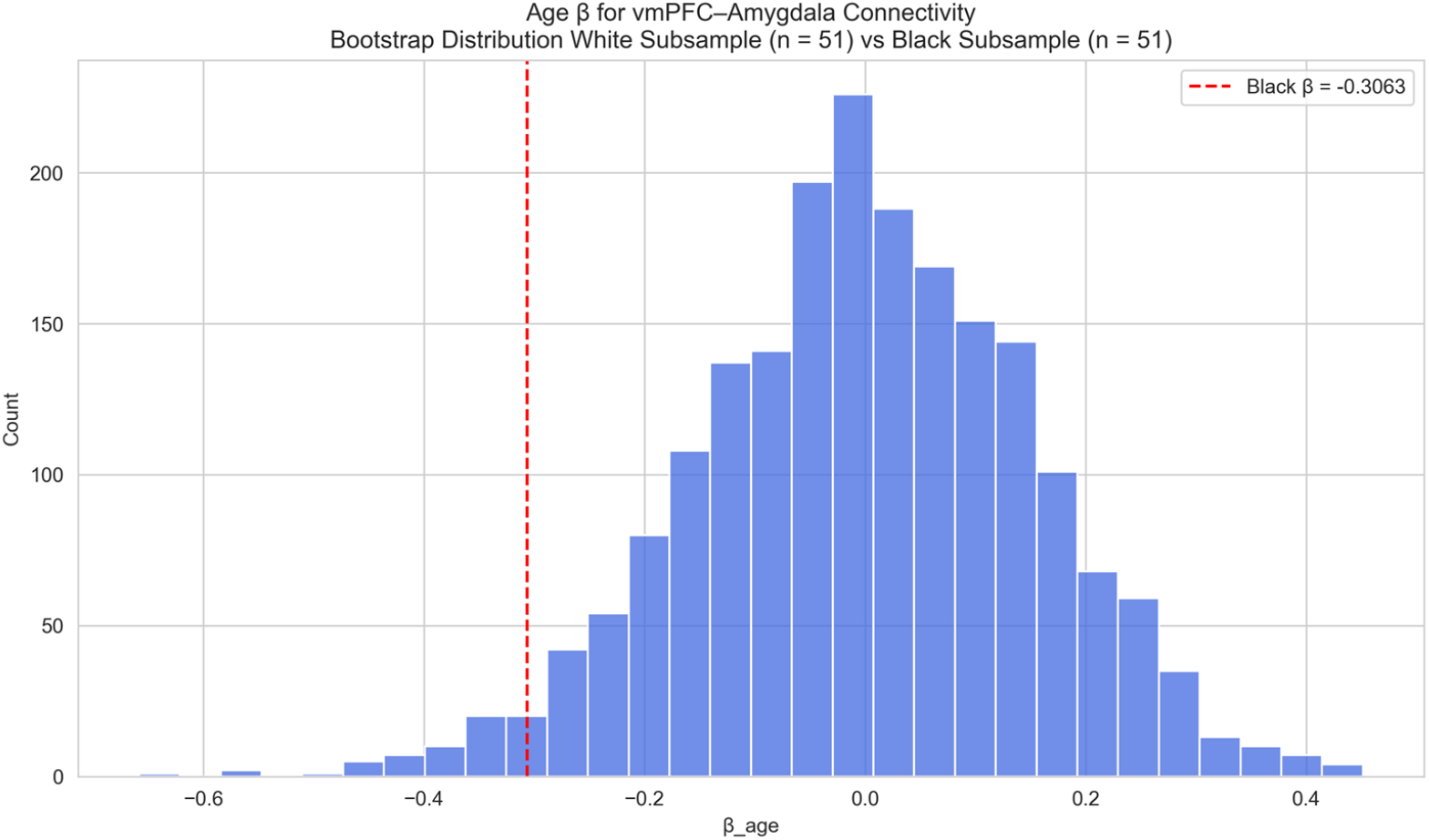
vmPFC–Amygdala Connectivity – Race-Stratified (Black vs White) Race-stratified analysis of age effects on vmPFC–amygdala functional connectivity: Black versus White participants. The observed age-related coefficient for the Black subgroup is shown as a vertical red line. Black participants showed a significantly stronger negative age-related effect on *vmPFC–amygdala* connectivity compared to matched White bootstrap samples (one-sided p = 0.027).

**Figure 11:**
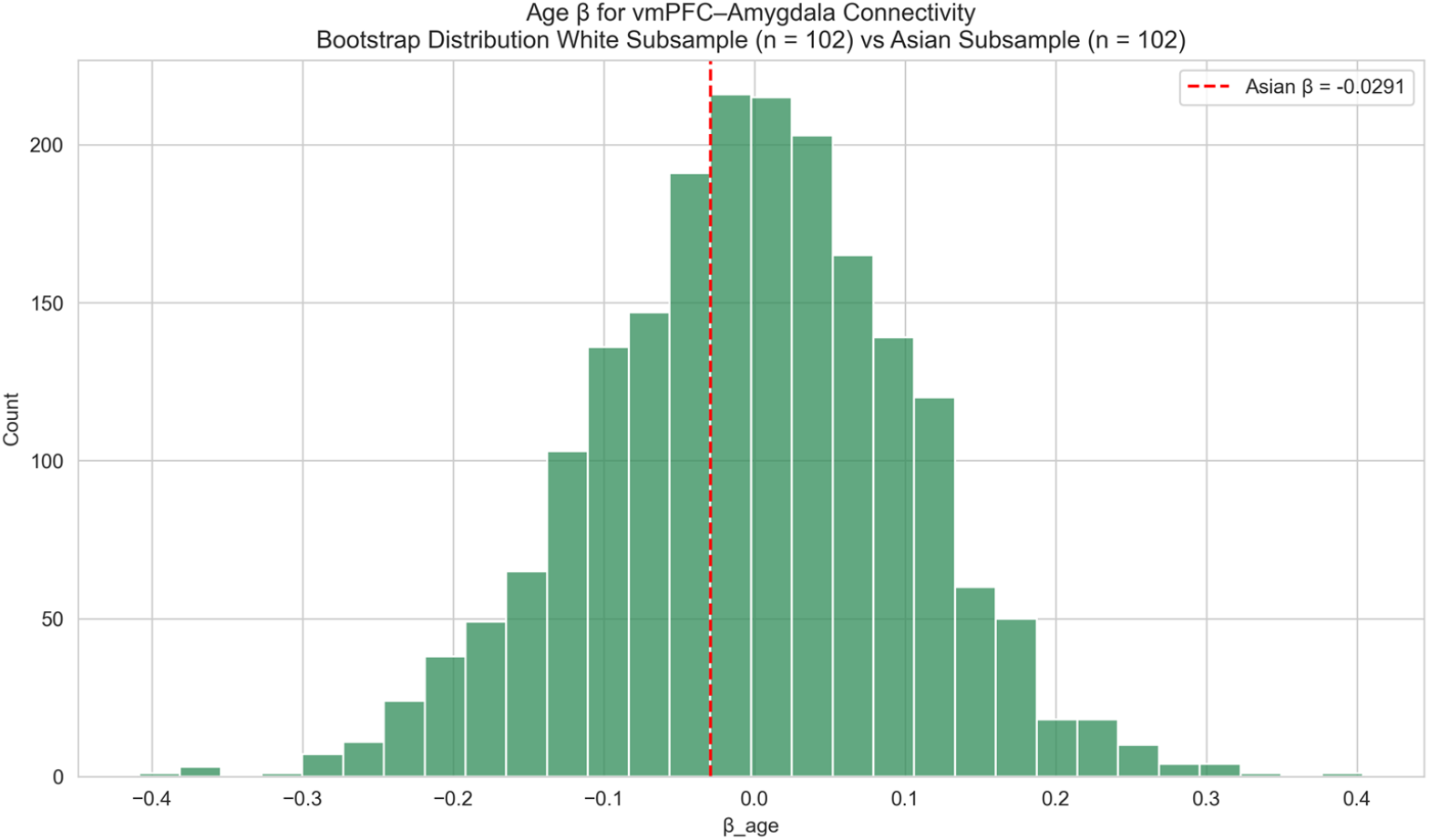
vmPFC–Amygdala Connectivity – Race-Stratified (Asian vs White) Race-stratified analysis of age effects on *vmPFC–amygdala* functional connectivity: Asian versus White participants. The observed age-related coefficient for the Asian subgroup is shown as a vertical red line. No significant age-related effects or differences were observed between Asian and White participants.

## Discussion

This study examined how age-related associations in emotion-related brain function, specifically amygdala activation and vmPFC–amygdala connectivity, may vary across race and SES using tfMRI data from the UK Biobank. Results revealed distinct patterns by social and demographic context. Amygdala activation was negatively associated with age in all racial groups. Additionally, participants with lower SES exhibited higher amygdala activation, independent of age. Notably, vmPFC–amygdala connectivity did not show significant associations with age or SES effects at the population level. However, the race-stratified model suggested a steeper age-related decline in connectivity among Black participants as compared to White participants.

These findings are broadly consistent with prior work suggesting that aging is associated with reduced amygdala reactivity, which may reflect more efficient emotional regulation in non-marginalized populations (Nashiro et al., 2012). Reduced amygdala activation in older adults has been linked to improved regulation of negative affect by increased vmPFC engagement, a pattern consistent with the “aging paradox” (Prakash et al., 2014). Although the overall sample revealed no age-related patterns in connectivity, stratified models suggested a steeper age-related decline in vmPFC–amygdala connectivity among Black participants as compared to White participants. These findings raise important questions about whether traditional neural models of aging may generalize across all demographic groups. Chronic exposure to structural stressors, such as racism and economic disadvantage, may shape neural aging differently in marginalized populations by intensifying allostatic load and cortisol production, which could lead to premature dysfunction of emotional regulation circuits (Fiocco, 2024).

The observed SES effect, where greater deprivation was linked to higher amygdala activation, is consistent with previous evidence that socioeconomic disadvantage is associated with heightened amygdala activity, potentially due to increased stress with economic instability (Migeot et al., 2022; Sinha et al., 2016). While SES did not moderate age-related associations in functional connectivity, this association with amygdala activation underscores the importance of examining brain-behavior relationships within the broader context of social determinants of health. Further, considering that the United Kingdom has lower racial diversity and a lower degree of income inequality than places like the United States (Dorling, 2015), the Townsend Deprivation Index may have limited sensitivity to thoroughly capture individual socioeconomic differences in this context (Trinidad et al., 2022).

This study is one of the few large-scale neuroimaging studies to examine racial and socioeconomic differences in age-related amygdala and vmPFC associations, underscoring the need for incorporating diversity, equity, and inclusion into neuroimaging research. Despite this, several limitations should be acknowledged. The number of Black and Asian participants with imaging data in the UK Biobank was disproportionately low compared to the White participants, limiting statistical power, generalizability, and representation. Although we used bootstrapping to improve robustness, larger and more representative datasets are needed to confirm these patterns in more detail. Additionally, race was self-reported, which may not fully capture the lived experiences of racism, discrimination, and cultural context that shape brain-behavior associations. Biological markers of allostatic load (e.g, telomere length, cortisol levels) (Murkey et al., 2022) could help clarify these effects.

Furthermore, although the amygdala and vmPFC are central emotion-related brain regions, they may not fully capture the complexity of emotional regulation networks. Incorporating additional regions and alternative functional tasks could provide a more comprehensive understanding. It is important to note that the Hariri Faces Emotions Task used in this study has demonstrated poor test-retest reliability, raising potential concerns about its effectiveness in detecting individual differences in brain-behavior associations (Elliott et al., 2020; Gee et al., 2015; Nord et al., 2017). Future studies would benefit from integrating multimodal imaging and psychosocial measures to more precisely model effects. These findings stress the need for inclusive brain health research that accounts for structural and social determinants across the lifespan.

## Acknowledgments

We are grateful to the UK Biobank and Oxford University for creating and producing the shared processed data. Computations were performed using the facilities of the Washington University Research Computing and Informatics Facility (RCIF). The RCIF has received funding from NIH S10 program grants: 1S10OD025200-01A1 and 1S10OD030477-01. Funding provided in part by the National Institutes of Health and Washington University in Saint Louis grant R25 NS130965-02.

## Notes

### Competing Interest Statement

The authors have declared no competing interest.

